# Generating gnotobiotic bivalves: a new method on Manila clam (*Ruditapes philippinarum*)

**DOI:** 10.1101/2024.05.08.593140

**Authors:** Marialaura Gallo, Andrea Quagliariello, Giulia Dalla Rovere, Federica Maietti, Barbara Cardazzo, Luca Peruzza, Luca Bargelloni, Maria Elena Martino

## Abstract

The microbiome, which encompasses microbial communities associated with animal hosts, exerts a profound impact on host physiology and ecosystem dynamics. The application of advanced sequencing technologies has enabled researchers to investigate the composition of microbiomes across a range of hosts and environments. While correlating microbial composition with health outcomes has been a priority, interpreting such data requires caution to avoid overemphasizing the roles of microbes. Understanding microbial influence demands mechanistic insights, which are often elucidated through gnotobiology. Despite their limitations in representing animal diversity, model organisms offer the advantage of reproducibility and experimental tractability. However, the marine realm, especially bivalves, which are crucial for ecosystem functioning and aquaculture, lacks gnotobiotic models. In this study, we present a method for generating microbiome-depleted and gnotobiotic clams (*Ruditapes philippinarum*), one of the most widely farmed molluscs in the world and a sentinel organism for climate change. This model expands gnotobiotic research into marine invertebrates, thereby enabling investigations into the impact of microbes on such key animal species.

## Introduction

The microbial communities associated with animal hosts, collectively known as the microbiome, are widely recognised for their significant influence on host physiology and the ecology of entire ecosystems [1,2]. As a result, understanding the composition and functions of animal microbiomes has received considerable attention over the past two decades. The ease of access to genomic data from many organisms, especially microbes, thanks to the rapid development of sequencing techniques, has resulted in a wealth of descriptive information, facilitating the understanding of microbiome composition and identity in different host environments, from plants and animals to broader ecosystems.

In this context, research on host-microbiome interactions has made extensive use of such data, correlating microbial composition with a variety of health and disease conditions [3–5]. However, the ease of data collection also presents a significant risk of obtaining information without a clear understanding of how to interpret them, potentially leading to an overemphasis on the role of microbiomes in animal health [6]. Identifying the mechanisms by which microbes shape specific host responses requires starting with a thorough description of microbial identity, ideally down to species or even strain level, and using carefully designed experimental methods aimed at establishing a causal relationship. In this respect, the long history of gnotobiology research, dating back to the late 19th century, has provided invaluable tools and insights. The generation of germ-free (GF) animals, ranging from guinea pigs to chickens, goats, and a variety of other mammals, birds, and amphibians [7–11], along with the creation of model organisms (i.e., *Drosophila melanogaster* [12], zebrafish [13], laboratory mice [14]) harbouring defined microbial communities, thanks to the development of microbiome transplant protocols, marked a revolutionary advance in host-microbiome research. This breakthrough has facilitated the testing of diverse hypotheses in different ecological contexts, with the aim of unravelling the intricate mechanisms by which microbes affect our lives.

Model organisms offer significant advantages, such as the high reproducibility of experiments across laboratories and the ease with which a variety of different ecological and molecular mechanisms can be dissected. However, they represent only a limited part of animal diversity and provide little or no information on species of major ecological or economic importance. In particular, the increasingly recognized importance of marine ecosystems calls for specific models to understand how host-associated microbial communities influence the ecology and evolution of marine animal hosts. The need for novel models is particularly relevant in the context of the rapid environmental modifications due to climate change. It is predicted that the effects of climate change will be particularly significant for coastal ecosystems, which are among the most productive and richest (in terms of biodiversity) habitats. Although most gnotobiotic research in aquatic animals has been conducted in zebrafish [13,15], successful efforts have been made to produce GF aquatic organisms beyond this species, such as the platyfish (*Xiphophorus maculatus*) [16], tilapia (*Tilapia macrocephala*) [11], several salmonid species [17], the sheepshead minnow (*Cyprinodon variegatus*) [18], and the starlet sea anemone (*Nematostella vectensis*) [19]. Most studies focused on vertebrate species, generally living in freshwater. Bivalve molluscs (e.g. clams, mussels and oysters) provide a crucial role in marine ecosystem functioning. They act as filter feeders, actively filtering water and particulates and creating substrates that serve as habitats for many other species [20]. One of the most recognised ecosystem services provided by bivalves is nutrient remediation. By filtering phytoplankton and accumulating nitrogen and phosphorus, they effectively absorb excess nutrients from the environment, including those from human activities such as agriculture and aquaculture [21]. Bivalves have also been proposed to act as carbon sinks or sources, but their exact contribution in this respect remains unclear. Because of their feeding behavior and their limited mobility, bivalves have been succefully used as bio-indicators of environmental quality [22]. Finally, extractive species, including bivalves, currently account for around half of all aquaculture production and have the potential to contribute significantly to the sustainable growth of the global aquatic food supply, although bivalve aquaculture will be the most affected by climate change [23].

Due to their ecological and economic importance, a gnotobiotic model for bivalves would therefore be essential to understand how host-associated microbial communities influence their response to environmental stressors and provide an experimental system to test hypotheses generated by the wealth of microbiome sequence data available for this taxonomic group.

In this study, we present a method for generating germ-free and gnotobiotic clams of the species *Ruditapes philippinarum*, the Manila clam. This is one of the most widely farmed molluscs in the world, it has been used in several studies on the effects of environmental pollution and its response to climate change has been assessed through an integrative biological approach [24].

The protocol presented here extends the current technical understanding of the establishment of gnotobiotic animals, providing the first model for a key taxonomic group in the marine realm. This will open new avenues for investigating the influence of microorganisms on animal health and elucidate the transferability of mechanisms predominantly studied in vertebrates to marine invertebrates.

## Materials and methods

### Clam maintenance and acclimation

Clams (average shell length 22.5 ± 1.9 mm, average soft tissue wet weight 1.0 g ± 0.2 g), were purchased from the SATMAR hatchery (France). A total of 90 clams were placed in a 20 L aquarium containing artificial seawater (ASW, Aquaforest Sea Salt) at 33 PSU salinity, with air stones to maintain constant water oxygenation and kept at room temperature. After 1 hour, 9 clams (3 per sample, 3 samples in total, T0, Table 1) were sacrificed for microbial load and 16S rRNA analysis. The aquarium was then placed in an incubator at an initial temperature of 18°C. To avoid acute thermal shock, the clams were acclimated for 5 days, during which the temperature was slowly increased up to 25°C. The final acclimation temperature (25°C) reflected the temperature of the subsequent experimental phases. The animals were fed once per day with New Coral Fito Concentrate (A.G.P., Italy), a commercial mixture of microalgae composed of Isochrysis (T-Iso) (33.3%) + Nannochloropsis (31 %) + Tetraselmis (18%) + Phaeodactylum (18%), at a final concentration of ∼40*10^6^ cells/L. Approximately 50% of the aquarium water was replaced twice during the acclimation period with fresh ASW at the appropriate temperature.

**Table 1.** The time points of clam sampling and experimental conditions (ATB: antibiotic).

| Time | Description |
| --- | --- |
| T0 | Received from hatchery |
| T1 | 5 days post-acclimation |
| T2 | 6 h post-ATB administration |
| T3 | 20 h post-ATB administration |
| T4 | 1 h post-transplant |
| T5 | 6 h post-transplant |
| T6 | 22 h post-transplant |

### Antibiotic treatment for microbial depletion in clams

#### Physical space considerations for optimal sterility

Prior to antibiotic treatment: *i)* all items, such as air stones and water pumps, were cleaned with 70% ethanol and placed under UV light for 20 minutes; *ii)* surfaces and equipment, such as incubators and aquaria, were cleaned with 70 % ethanol; and *iii)* the artificial seawater was plated on rich microbiological media such as Marine Agar (MA) and Luria Bertani (LB) agar (Condalab, Spain) to check its sterility.

#### Antibiotic treatment

After 5 days of acclimation, 9 clams (3 per sample, 3 samples in total, T1, Table 1) were sacrificed for microbial load and 16S rRNA analysis. The remaining clams (n=60) were divided into four aquaria. Each aquarium was set up with 15 clams and 3 L of ASW and kept at 25°C. Three aquaria were used for antibiotic treatment and the remaining one was used as a control group (i.e. no antibiotic treatment) and placed in a separate incubator. Treated groups received a mixture of five antibiotics at two times: T1, i.e. at the end of the acclimation period and T2, i.e. 6 h after the first antibiotics administration.

The antibiotics were administered by pipetting the mixed solutions into the treatment tanks. To increase the uptake of the antibiotics, the doses were administered after the clams had been fed. The antibiotic mixture consisted of erythromycin (83 mg/L), ampicillin (83 mg/L), streptomycin sulphate (20 mg/L), ciprofloxacin (20 mg/L) and cefotaxime sodium (20 mg/L). Erythromycin and streptomycin act by inhibiting bacterial protein synthesis [25,26], ampicillin and cefotaxime target bacterial cell wall synthesis [27], while ciprofloxacin targets nucleic acid synthesis [28]. Stock solutions for each antibiotic compound were prepared freshly in Milli-Q water or 20% ethanol prior to each administration according to the manufacturer’s instructions. The control group received an equivalent volume of Milli-Q water and 20% ethanol. The set of antibiotics and corresponding concentrations were chosen to maximise bacterial perturbation and were validated in preliminary experiments (data not shown). At T2 and T3 (6 h and 20 h after antibiotic administration, respectively, Table 1), three clams from each aquarium were collected, sacrificed and pooled (1 sample per aquarium) for assessment of microbial load and 16S rRNA analysis.

### Microbiome transplant

#### Selection of the mock community

The bacterial strains used to generate the mock community for microbiome transplant were isolated from clam homogenate. Six clams taken from the acclimation aquarium were crushed together for 2 min at maximum speed using the Stomacher® 3500 instrument (VWR, Italy). Subsequently, 100 µl of the homogenate were plated on different selective and/or differential media, including MA, Iron agar and Thiosolfate Citrate Bile Sucrose (TCBS) agar (Condalab, Spain), after appropriate serial dilution. The plated samples were then incubated at 22°C for 48-72 hours. Colonies were selected on the basis of morphological differences and part of them was used for DNA extraction. The remaining colonies were subcultured on MA at 22°C for 48-72 hours. After incubation, cultured samples were collected and stored in 80% glycerol at −80°C.

For DNA extraction, the colony was dissolved in 100 µl Milli-Q water and boiled at 95°C for 10 minutes. After centrifugation at 10,000 rpm for 5 min, the pellet was discarded and the DNA-containing supernatant was immediately used for 16S rRNA gene amplification (see section below for details on 16S rRNA amplification) as described previously. The resulting PCR products were visualised by 1.5% agarose gel electrophoresis and subsequently purified using the ExoSAP™ PCR Product Cleanup Kit (Applied Biosystems, USA). Sanger sequencing of the 16S rRNA gene was performed at BMR Genomics Company, Italy. Species identification of each colony was achieved by comparing the partial 16S rRNA gene sequences obtained (100 to 450 nt) with those in the GeneBank database, using web-based BLAST software [29]. To obtain the bacterial suspension to be used for microbiome transplant, each bacterial colony was inoculated into 50 ml of Marine Broth (Condalab, Spain) and cultured at 22°C for up to 48 h to reach 10^8^ CFU/ml.

#### Transplant of the mock community

For the transplant procedures, a total of 90 clams purchased from the SATMAR hatchery (France) were acclimated under the same conditions as described above. After acclimation, clams were divided into four aquaria (15 clams per aquarium). Germ-free (GF) clams were obtained for all aquaria as described above. 20 h after antibiotic administration (T3, Table 1), GF clams were transferred to small aquaria containing 1.5 L of ASW for the 2 hour depuration phase. Subsequently, 12 GF clams from individual tanks were transferred to new tanks containing 450 ml of ASW. To produce gnotobiotic clams, 50 ml of bacterial suspension containing 10^8^ CFU/ml *Vibrio diazotrophicus*, 10^8^ CFU/ml *Shewanella colwelliana* and 10^8^ CFU/ml *Halomonas alkaliphila* was added directly to two tanks. The remaining two tanks were used as controls and 50 ml of ASW was added. One hour after bacterial inoculation, 3 clams from each tank (transplanted and controls) were collected, sacrificed and pooled (1 sample per tank) for assessment of microbial load and 16S rRNA analysis (T4, Table 1). The contents of each tank were transferred to a new aquarium containing 2.5 L of ASW (a total of 3 L of ASW per aquarium with 9 transplanted or control clams) and kept at 25 °C. After 6 h (T5, Table 1) and 22 h (T6, Table 1), 3 clams from each tank were collected, sacrificed and pooled (1 sample per tank) for microbial load assessment and 16S rRNA analysis.

### Clam collection and dissection for microbial count and 16S rRNA analysis

The clams were collected at the time points listed in Table 1:

For T0 and T1, three samples of 3 clams each were collected from the same aquarium and processed separately. For T2 and T3, three clams were collected from each aquarium (3 treatment and 1 control – 4 samples in total), sacrificed and pooled as follows. For each clam, the adductor muscle was cut, the valves were opened and the entire wet body was collected in a stomach bag together with the extrapallial fluid using sterile tweezers and a pipette. Three clams were collected in the same stomacher bag and homogenised together by adding 9 volumes of phosphate buffered saline (PBS) (Sigma-Aldrich, Germany) of the sample weight for 2 minutes at maximum speed using the Stomacher® 3500 (VWR, Italy). To assess the microbial load, samples were serially diluted in PBS, plated on MA (Condalab, Spain) and incubated at 22°C for 48h. For subsequent RNA extraction, 2 ml of the homogenised samples were centrifuged at 4000 rpm for 2 min, and the pellet was stored at -80°C.

### RNA extraction, amplification and sequencing

For RNA extraction, 0.5 µm glass beads were added to the previously stored pellet, which was then processed using the RNeasy Mini Kit (Qiagen, Germany) according to the manufacturer’s instructions. RNA concentration was measured using a NanoDrop ND-1000 spectrophotometer (Thermo Scientific, USA). For microbiota characterisation, 1 µg of RNA was reverse transcribed into cDNA using the SuperScript™ IV First-Strand Synthesis System (Invitrogen™, USA).

The 16S rRNA gene was amplified using two universal 16S primers (forward primer, UniF 5′-GTGSTGCAYG GYTGTCGTCA-3′ and reverse primer, UniR 5′-ACGTCRTCCMCACCTTCCTC-3′ [30]. The 16S rRNA gene of *Endozoicomonas* spp. was amplified using an *Endozoicomonas*-specific primer set including a reverse primer (En771R: 5′-TCAGTGTCARRCCTGAGTGT-3′) and a bacterial universal forward primer (27F: 5′-AGAGTTGATCMTGGCTCAG-3′) [31]. End-point PCRs were performed in a total of 20 µl on Mastercycler® nexus SX1 (Eppendorf, Hamburg) using DreamTaq™ PCR Master Mix (Thermo Scientific, USA). Reaction mixtures consisted of 0.5 µl of each primer, 10 µl PCR Master Mix, 7 µl water and 2 µl DNA (or cDNA) template. PCR conditions included 1 cycle of initial denaturation at 94 °C for 2 min, followed by 35 cycles of denaturation at 94 °C for 20 s, annealing at 56 °C for 30 s, extension at 72 °C for 30 s and a final extension at 72 °C for 7 min. PCR products were observed by 1.5% agarose gel electrophoresis.

For 16S rRNA gene amplicon sequencing, library preparation and sequencing of the V3–V4 hypervariable regions of the bacterial 16S rRNA gene were performed at BMK Gene (Germany). Sequencing was performed on an Illumina Novaseq 6000.

### 16S rRNA sequencing analysis

Raw data were processed in order to remove adapters, select reads lengths and remove low-quality reads using Fastp [32], Trimmomatic v0.33 [33] and cutadapt 2.7.8 [34]. Cleaned reads were then processed for downstream analyses using R software with dada2 [35] to identify Amplicon Sequence Variants (ASV), and obtain a taxonomic identification using silva_nr99_v138.1_wSpecies_train_set.fa as database. Biodiversity metrics were estimated using *phyloseq* [36] and *vegan* packages in R, with Bray-Curtis distance used for Beta-Diversity.

## Results

### Generation of germ-free clams

In this study, we developed a protocol for the microbiological sterilisation of adult Manila clams by administering a mixture of antibiotics in the water (Fig. 1). After five days of acclimation at 25°C (T1), the clams were divided into four tanks: three of which were treated with a mixture of five antibiotics, while the remaining tank was used as a control. The four tanks were kept at 25 °C for 24 hours.

**Fig 1.**
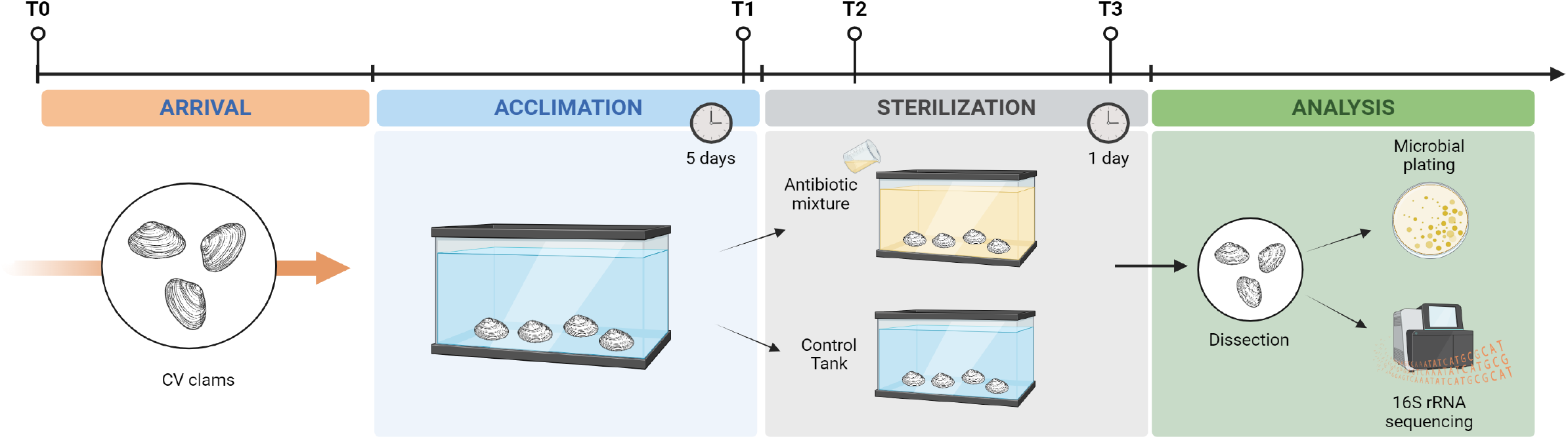
Schematic diagram of the experimental protocol for the production of germ-free clams (CV: conventional clams; T0: arrival; T1: 5 days post-acclimation; T2: 6h post antibiotic administration; T3: 20h post antibiotic administration).

To assess the efficacy of the antibiotic treatment, we first monitored microbial growth from T0 (when the clams were purchased from the hatchery) to T3 (after two doses of antibiotics) by plating the clam homogenate on Marine Agar (MA) culture medium, a non-selective medium commonly used to grow and isolate a wide variety of heterotrophic marine bacteria. Starting from a total microbial load of 10^6^ CFU/ml (T0 - n=3 clams), microbial growth decreased over time and no bacterial growth was detected after two doses of antibiotics. One dose of antibiotics resulted in a two-log reduction in microbial load (T1, Fig. 2A). To assess the presence of viable but nonculturable bacteria in the treated clams, we also performed 16S rRNA sequencing of RNA extracted from the clam homogenate. A total of seven samples were sequenced on Illumina Novaseq6000 sequencing platform, generating 8,527,652 pair raw reads. These PE reads were processed for quality control, assembly and data filtration, which yielded 6,402,125 clean reads. A minimum of 50,186 clean reads were generated for each sample and the average data output per sample was 72,751 clean reads. Taxonomic analyses at the genus level revealed a significant decrease in the microbial community of clams following antibiotic treatment, while control clams (i.e., not treated with antibiotics) showed an increase in the number and type of bacterial genera (Fig. 2B, Table S1). After acclimation (T1), the clam microbiome was characterised by a total of 26 bacterial genera, with the predominant presence of *Endozoicomonas* (82.9%), followed by *Pseudomonas* spp. (4.1%), *Bacillus* spp. (2.5%), *Umboniibacter* spp. (2%), *Salinirepens* spp. (1.7%), *Oleiphilus* spp. (1.3%), *Stenotrophomonas* spp. (1.2%) and 19 other bacterial genera, representing less than 1% each (Table S1, Fig. 2B). The administration of the antibiotic mixture caused a significant decrease in the diversity of the clam microbial community (Fig. S1), which was dominated by *Endozoicomonas* spp. (98%) (Fig. 2B). The abundance of *Bacillus* spp. decreased to 1% and traces of five other bacterial genera were also detected, but their abundance was less than 0.01% (Table S1, Fig. 2B). However, the prevalence of 24 bacterial genera was observed in the control aquarium. In particular, the dominance of the genus Endozoicomonas, which was prominent at the end of acclimation and resistant to antibiotic treatment, decreased to 46% in untreated clams. This reduction was accompanied by an enrichment of other bacterial genera, including *Malaciobacter* spp. (8%), *Umboniibacter* spp. (7.4%), *Neptuniibacter* spp. (6%), *Alteromonas* spp. (6%), *Pontibacterium* spp. (5%) *Salinirepens* spp. (4.5%), *Bacillus* spp. (2%), Marinomonas spp. (2%), *Aliiroseovarius* spp. (1.7%), *Marinobacter* spp. (1.4%), *Lentibacter* spp. (1.3%), *Acinetobacter* spp. (1%), *Sulfitobacter* spp. (1%) and 10 other bacterial genera, each representing less than 1% (Table S1, Fig. 2B).

**Fig 2.**
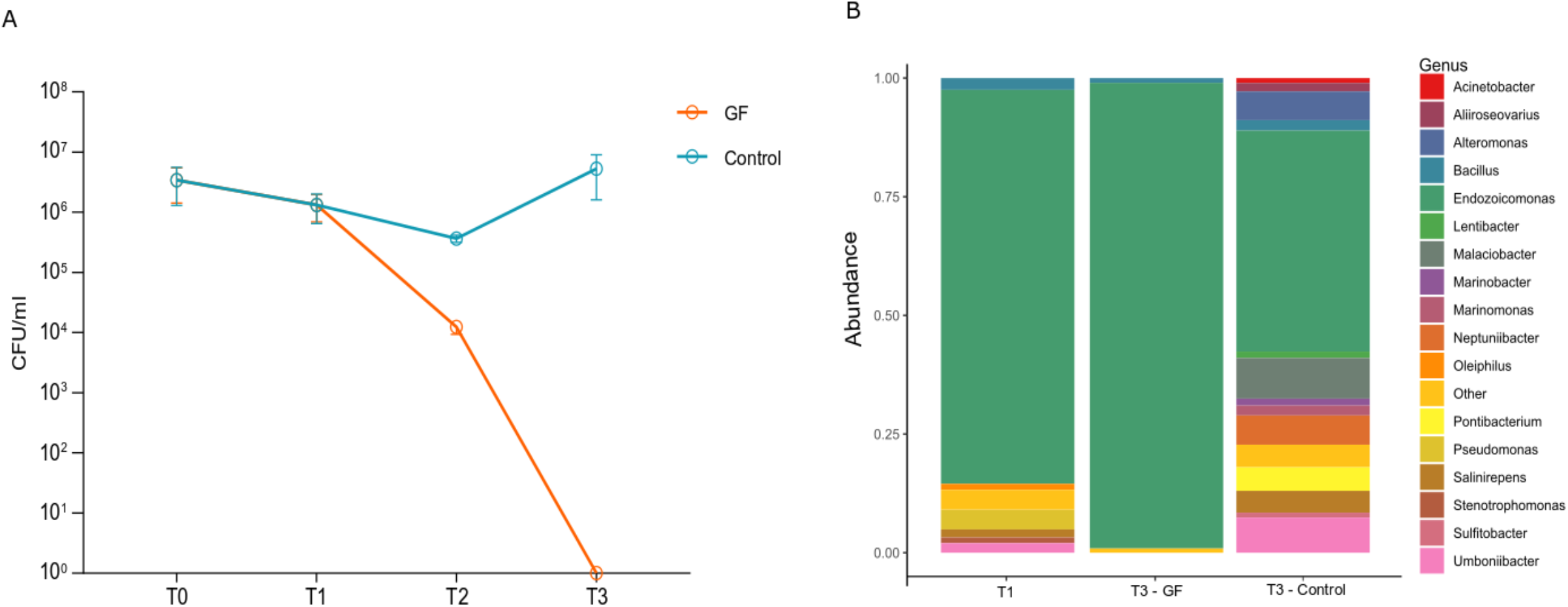
(**A**) Microbial load (CFU/ml) detected in clams after antibiotic treatment (GF) and in control animals (T0: arrival; T1: 5 days post-acclimation; T2: 6h post antibiotic administration; T3: 20h post antibiotic administration). (**B**) Taxonomic composition of relative microbiome abundance at genus level of acclimated clams (T1), antibiotic treated (T3 - GF) and control clams (no antibiotic treatment; T3 - Control). Bar plot of significant genera (genera with relative abundance <0.1 % are grouped as “Other”).

Overall, our results demonstrate that the protocol developed to obtain germ-free clams resulted in the near-complete depletion of the clam microbiome, with almost exclusively *Endozoicomonas* spp. persisting in the antibiotic-treated clams. To classify the antibiotic-resistant *Endozoicomonas* at the species level, we Sanger sequenced the 16S rRNA of the treated clams using primers specific for *Endozoicomonas* spp.. The resulting sequence was identified as *Endozoicomonas elysicola* (File S2).

### Generation of gnotobiotic clams

#### Selection of bacterial species for the mock community

In order to carry out the transplant experiment, we sought to create a mock community consisting of known concentrations of defined bacterial species, capable of colonising bivalve molluscs. To achieve this, six acclimated clams (T1) were collectively crushed and the resulting homogenate was plated on selective and differential media. Following incubation, three colonies were randomly selected based on their unique morphology on MA for 16S rRNA gene sequencing for species-level classification. The species were finally identified as *Vibrio diazotrophicus*, which is characterised by small round, smooth, opaque white colonies with a halo, *Shewanella colwelliana*, showing round, smooth, pink colonies, and *Halomonas alkaliphila*, which has round, smooth, convex and creamy white colonies.

#### Microbiome transplant

To proceed with the transplant of the mock community, we first treated the Manila clams with the antibiotic regimen as described above (Fig. 1). Twenty hours after antibiotic administration (T3), clams were depurated from antibiotics for two hours before being transferred to small tanks with fresh ASW (Fig. 3). The microbial mock suspension, consisting of the three selected bacterial species (final volume: 50 ml; [C] = 10^8^ CFU/ml per species), was added directly to the water containing the clams. The control tanks received an equal volume of ASW (Fig. 3). After one hour of incubation (T4), the entire contents of the small tanks (including clams and water, with or without bacterial suspension) were transferred to new tanks containing fresh ASW (total volume = 3 L per tank).

**Fig 3.**
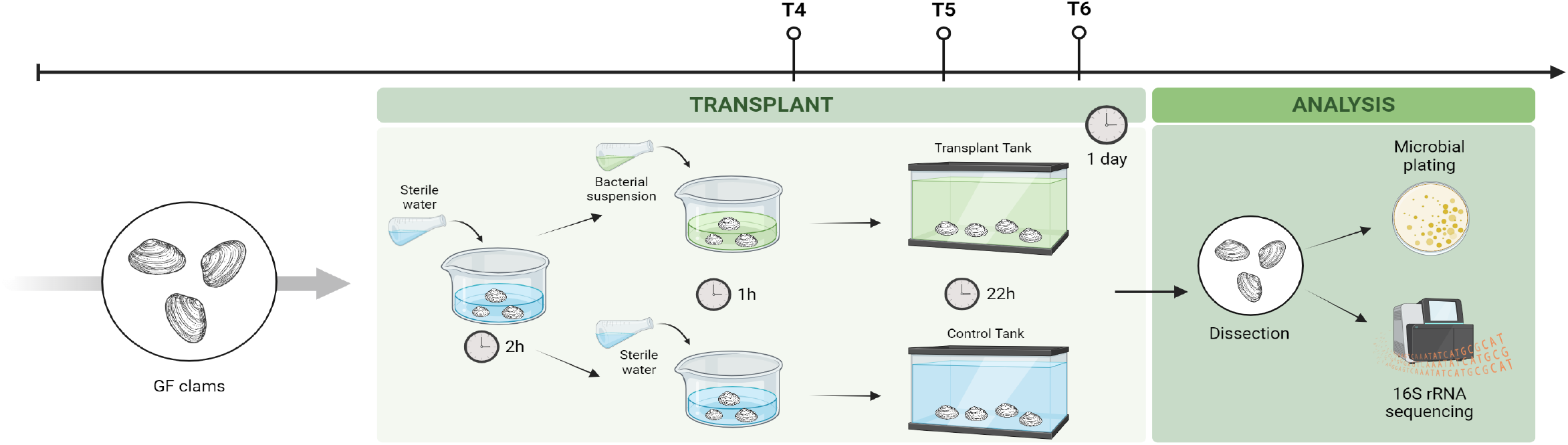
Schematic diagram of the experimental protocol for the microbiota transplant in germ-free (GF) clams (T4: 1h post transplant; T5: 6h post transplant; T6: 22h post transplant).

To assess the efficacy of the microbiome transplant, we monitored bacterial growth throughout the experiment (Fig. 4A). No bacterial growth was detected after antibiotic treatment (T4, Fig 4A). In the transplanted clams, a rapid and significant increase in microbial load was observed as early as one hour after transplant (T4 - 10^4^ CFU/ml), with a final concentration of 10^6^-10^5^ CFU/ml (T5 and T6, respectively). In contrast, control clams (i.e. animals that received antibiotic treatment but no microbiome transplant) showed a detectable microbial load at 8 hours after transplant (T5 - 10^2^ CFU/ml), reaching a final concentration of 10^6^ CFU/ml at T6 (Fig. 4a).

**Fig 4.**
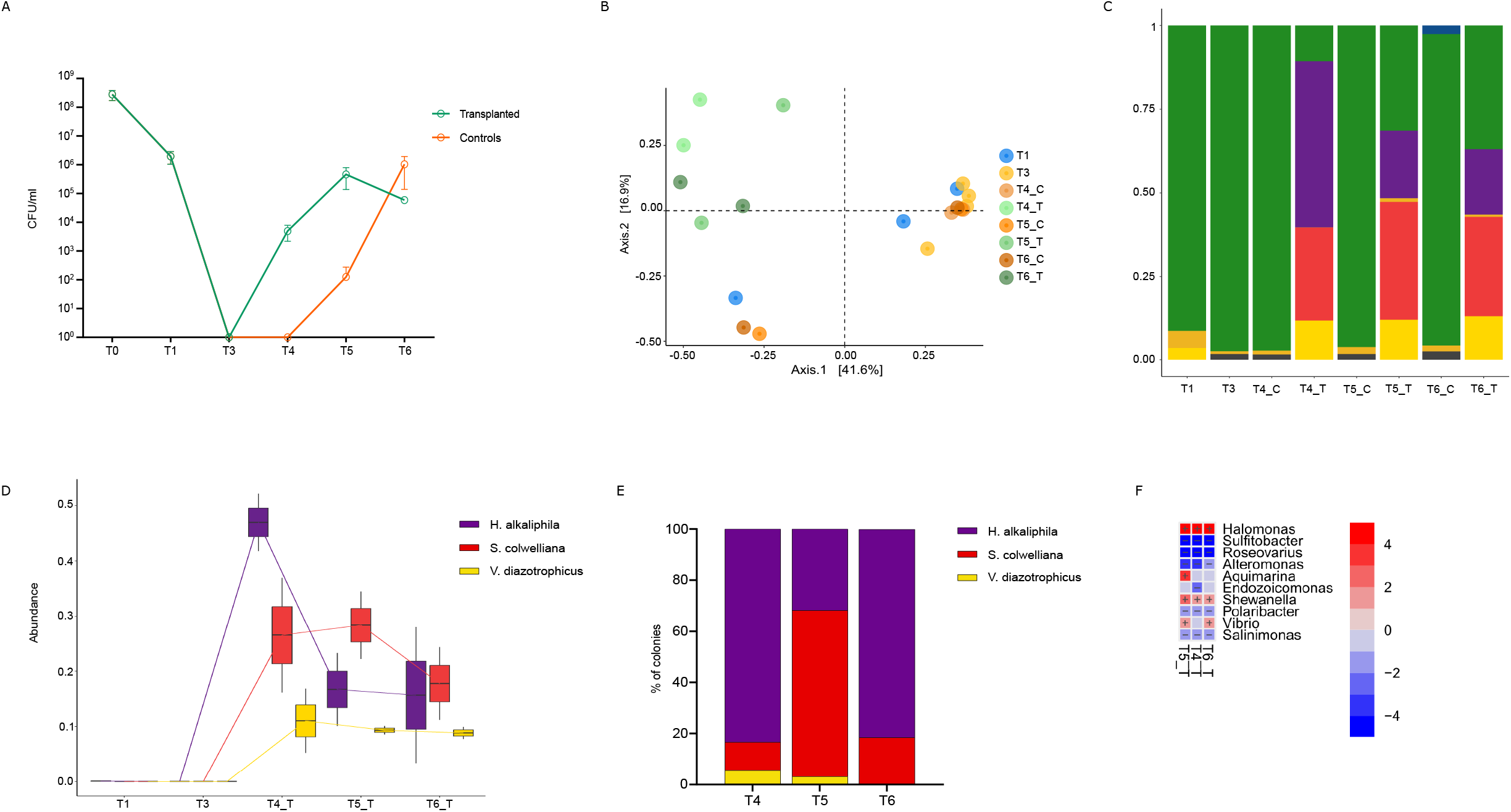
(**A**) Microbial load (CFU/ml) detected in clams throughout the experiment. The description of the time points is given in Table 1. (**B**) PCoA (at ASV level) illustrating sample clustering based on bacterial treatment and time (C: control clams; T: transplanted clams). (**C**) Taxonomic composition of relative microbiome abundance at genus level. Genera with relative abundance <1 % are grouped as “Other”. d, e) Relative abundance of the three bacterial species selected for microbial transplant obtained as determined by 16S rRNA sequencing (**D**) and microbial plating on MA (**E**). (**F**) Maaslin2 results on transplanted clams using T3 as fixed effect.

To determine the effect of microbial transplant on the composition of the bacterial community, we performed 16S rRNA amplicon sequencing of the transplanted and control individuals. A total of 22 samples were sequenced on Illumina Novaseq6000 sequencing platform, generating 10,943,861 pair reads. A minimum of 220,468 clean reads were generated for each sample and the average data output per sample was 497,448 clean reads. Principal Coordinates Analysis (PCoA) revealed a clear clustering between transplanted and control individuals (Principal Component 2 - PC2; Fig. 4b). This suggests that a differentiation of the clam microbiome has occurred following microbial transplant. While the control samples (i.e., non-transplanted clams) maintain a stable association with *Endozoicomonas* spp. (Fig. 4c), the bacteria that contribute to the post-transplant differentiation are indeed the three inoculated bacterial species that were detected and significantly increased only in the transplanted clams, both by 16S rRNA amplicon sequencing and by plating (Fig. 4d-e, Table S3). In particular, *H. alkaliphila* and *S. colwelliana* colonized the transplanted individuals more efficiently, while *V. diazotrophicus* was recovered at lower levels (Fig. 4d-e, Table 2). Interestingly, all transplanted species showed a significant increase in recipient clams, which was accompanied by a significant decrease in six bacterial species, suggesting outcompetition by the transplanted species (Fig. 4f Table S3).

**Table 2.** Relative abundance of the three bacterial species selected for microbial transplant obtained as determined by 16S rRNA sequencing and microbial plating of transplanted clams.

| Transplanted Species | Relative Abundance |  |  |  |  |  |
| --- | --- | --- | --- | --- | --- | --- |
|  | 16S amplicon sequencing |  |  | Microbiological plating |  |  |
|  | T4 | T5 | T6 | T4 | T5 | T6 |
| *Halomonas alkaliphila* | 49.6 | 20.1 | 19.5 | 83.4 | 31.8 | 81.6 |
| *Shewanella colwelliana* | 27.8 | 35.2 | 19.529.6 | 11.1 | 65.1 | 18.4 |
| *Vibrio diazotrophicus* | 11.8 | 12.1 | 13.1 | 5.5 | 3.1 | 0 |

## DISCUSSION

The generation of gnotobiotic animals represents a powerful strategy to study the mechanisms underlying the relationship between animal hosts and their microbiome at different levels, allowing the dissection of physiological response down to molecular processes. Since the late 19^th^ century, various animals have been sterilised for research purposes. During this time, basic research on host-microbe interactions has been conducted with germ-free and gnotobiotic model organisms, from invertebrates (*C. elegans, Drosophila melanogaster*, etc.) and vertebrates such as zebrafish and mice. However, the nature of host-microbe relationships varies greatly across hosts and ecosystems, as many of them are highly dependent on the environment. Therefore, it is essential to study dynamics and processes in non-model organisms and complex systems. Here, we present a new method to generate gnotobiotic bivalves using the Manila clam *Ruditapes philippinarum* as a model species. This species, which is found in lagoons and river deltas, is of great importance from an environmental standpoint, as it provides a range of ecosystem services [21]. Additionally, it is a valuable economic resource for global aquaculture, offering benefits (e.g. economic and societal) to local communities of producers, which are often centered around bivalve farms.

In this work, we have developed a protocol for microbiome depletion and transplant on adult clams. The microbiome of the treated clams consisted mainly of *Endozoicomonas* spp., followed by *Pseudomonas, Umbonibacter* and a few other bacterial genera (Fig. 2b). The microbiome-depletion protocol successfully reduced all bacterial genera, with the exception of *Endozoicomonas* spp., which was only detected by 16S rRNA amplicon sequencing and consequently identified as *Endozoicomonas elysicola* by Sanger sequencing. No bacterial growth was observed after 20 hours of antibiotic treatment, indicating that *E. elysicola* was not culturable on MA. This is noteworthy as *Endozoicomonas* spp. have been shown to be cultivable on general media such as MA [37,38], suggesting that the strain present in the treated clams may have specific nutritional requirements and/or be difficult to isolate. *Endozoicomonas* spp. are prevalent symbionts in a variety of marine hosts, including corals [39], and other cnidarians [40], sponges [41], gorgonians [42], worms [43], fish [44], tunicates [45] and molluscs [46]. They generally reside in aggregates within the host endodermal tissues [39]. For these reasons, it has not always been easy to isolate *Endozoicomonas* from host tissues [47] and despite its associations with numerous hosts in oceans worldwide, the functional role of *Endozoicomonas* remains unclear. Further studies are needed to investigate where *Endozoicomonas* resides (i.e., in organs or tissues) using imaging techniques, which have been shown to be effective in revealing the spatial distribution of such species [38,39]. This would be important information for the development of targeted treatments to effectively deplete the species.

The protocol we developed for microbiome transplantation on adult Manila clams is based on the inoculation of a mock community consisting of three bacterial species. The method proved to be effective in transplanting the three bacterial species, although the final abundances in the recipient clams differed. Indeed, the recipient animals showed a distinct microbiome profile in comparison to the control animals (Fig. 4b), with a significant increase in the abundance only of the transplanted species (Fig. 4c,d). This result was also confirmed by classical microbiological plating (Fig. 4e). Transplantation appears to be effective as early as one hour after inoculation of the bacterial communities and the species were still found after 20 hours. Notably, a single microbiome transplantation appears to lead to a slight decrease of the transplanted species over time, with the exception of *V. diazotrophicus*, which appears to efficiently colonize the clams and be stable over time (Fig. 4c, d, f). It would be interesting to test the effect of multiple transplantations on recipient clams to further extend the efficacy of transplantation, as well as to test the efficiency of the protocol in the longer term. It is also noteworthy that although the antibiotic treatment was effective for most bacterial symbionts of the clams, with the exception of *E. elysicola*, the transplantation method allowed a significant reduction in the relative abundance of this species, with all three transplanted species contributing to its outcompetition (Fig. 4c). These results are particularly intriguing in the context of the need to reduce antibiotic use, including in aquaculture systems. They demonstrate that harnessing the higher fitness of symbionts, especially beneficial symbionts, could be a more effective strategy for outcompeting pathogens. This paves the way for targeted treatments based on the administration of probiotics and beneficial symbionts to animals in aquaculture facilities.

Although our technique is highly effective in significantly reducing the microbiome of clams and generating gnotobiotic animals, technical challenges need to be addressed to further advance the study of gnotobiotic bivalves. First, improved methods to efficiently isolate the aquaria (e.g., seals, sterilized pumps, etc.), as well as automated water filtration methods need to be developed to completely eliminate environmental contamination. Secondly, testing our protocols on clams of different origins, possibly carrying different microbiomes, would allow us to test the applicability of our techniques under different ecological conditions. Finally, the definition of a sterile diet could represent an added value allowing different laboratories conducting microbiome research on bivalves to standardize the nutritional regimes.

In conclusion, this newly developed technique to generate gnotobiotic bivalves represents a significant advance in host-microbiome research, particularly in studies seeking to identify causal effects. This will help to further strengthen the analysis of the impact of microbes on animal health in marine ecosystems, thereby expanding the potential of gnotobiotic research.

## Supporting information

Supplemental Figure 1. Observed and Shannon indexes were estimated to evaluate the alpha diversity among samples.

