## Supplementary figures and images for "Generating gnotobiotic bivalves: a new method on Manila clam (*Ruditapes philippinarum*)"

### Supplemental Figure 1. Observed and Shannon indexes were estimated to evaluate the alpha diversity among samples.

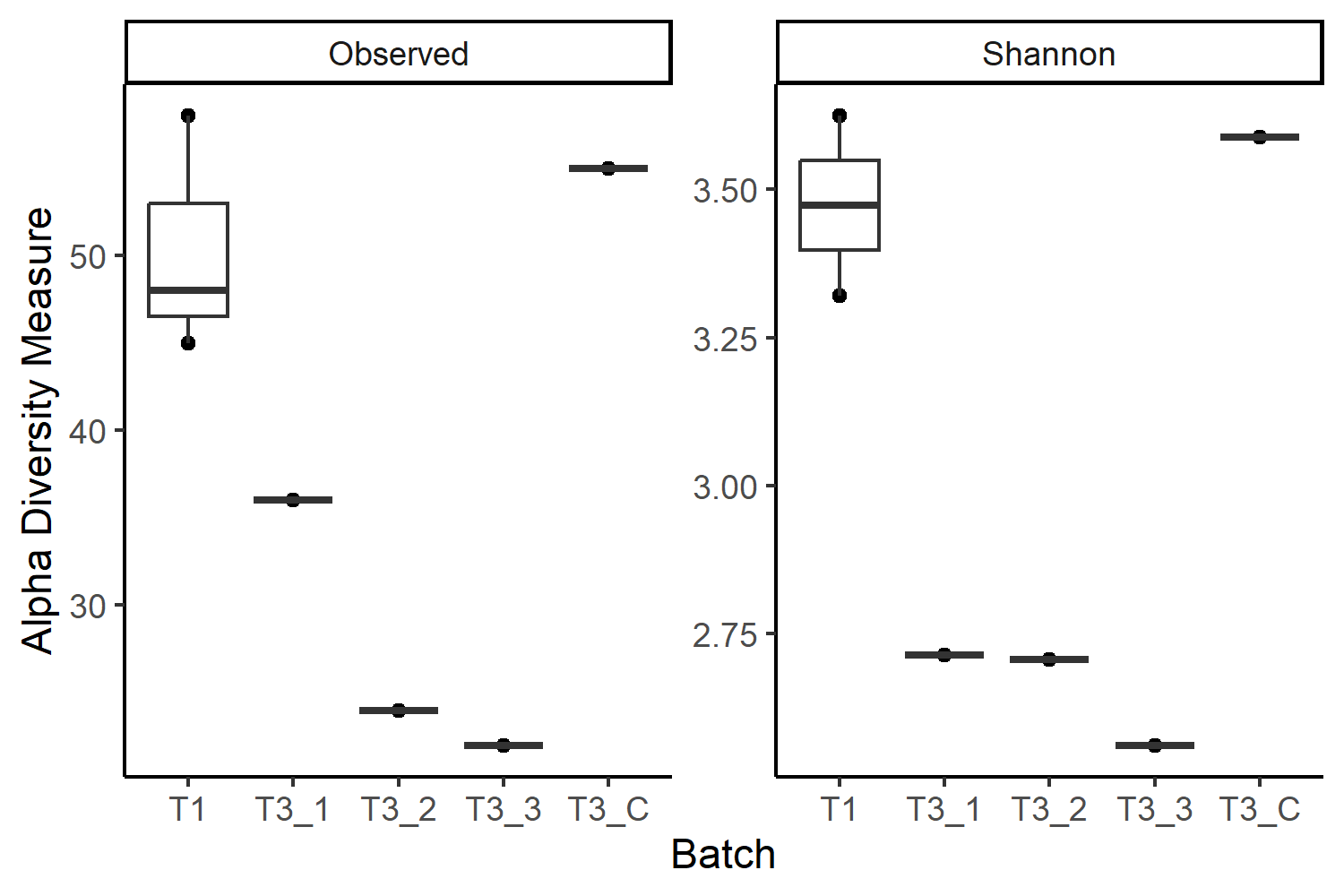
