## Supplemental Table 1 for "Generating gnotobiotic bivalves: a new method on Manila clam (*Ruditapes philippinarum*)"

**Table S1.** Relative abundances of all bacterial genera detected in the experimental samples through 16S rRNA sequencing (T1: acclimated clams; T3-GF: antibiotic-treated clams; T3-Control: conventional clams)

| **Genus** | **Abundance** | | |
| --- | --- | --- | --- |
|  | **T1** | **T3-GF** | **T3-Control** |
| Endozoicomonas | 0,829918781 | 0.98095748 | 0,4655532359 |
| Pseudomonas | 0,04184666681 | 0 | 0 |
| Bacillus | 0,02486915939 | 0.010149003 | 0,02192066806 |
| Umboniibacter | 0,02018231202 | 0.000340657 | 0,07411273486 |
| Salinirepens | 0,01676208074 | 0 | 0,04592901879 |
| Oleiphilus | 0,01262629933 | 0 | 0 |
| Stenotrophomonas | 0,01213049387 | 0 | 0 |
| Alcanivorax | 0,005030181087 | 0 | 0,007306889353 |
| Cerasicoccus | 0,005011933514 | 0 | 0 |
| Yoonia-Loktanella | 0,00393442623 | 0 | 0 |
| Sva0996 marine group | 0,003688799463 | 0.00194312 | 0 |
| BD1-7 clade | 0,003476749239 | 0 | 0 |
| Mycoplasma | 0,003440314015 | 0 | 0 |
| NS3a marine group | 0,003315853399 | 0 | 0 |
| Neptuniibacter | 0,003018108652 | 0 | 0,06158663883 |
| Brevundimonas | 0,00234741784 | 0 | 0 |
| SM1A02 | 0,001676727029 | 0 | 0,002087682672 |
| Truepera | 0,001006036217 | 0.000832271 | 0 |
| Marinobacterium | 0,0008743169399 | 0 | 0,01461377871 |
| Marinobacter | 0,0008691873099 | 0 | 0 |
| [Eubacterium] fissicatena group | 0,0008691873099 | 0 | 0 |
| Crocinitomix | 0,0006706908115 | 0 | 0 |
| Cutibacterium | 0,0006706908115 | 0 | 0 |
| Spongiibacter | 0,0006706908115 | 0 | 0 |
| Flavicella | 0,0006557377049 | 0.000321285 | 0 |
| Litoricola | 0,0004371584699 | 0 | 0 |
| Nitrosomonas | 0 | 0.001124498 | 0 |
| Nitrospira | 0 | 0.001021972 | 0 |
| Aureimarina | 0 | 0.000851644 | 0 |
| Amphritea | 0 | 0.000803213 | 0 |
| Paenibacillus | 0 | 0.000803213 | 0 |
| Ulvibacter | 0 | 0.000510986 | 0 |
| Acinetobacter | 0 | 0.000340657 | 0,01043841336 |
| Malaciobacter | 0 | 0 | 0,08559498956 |
| Alteromonas | 0 | 0 | 0,06054279749 |
| Pontibacterium | 0 | 0 | 0,05010438413 |
| Marinomonas | 0 | 0 | 0,02087682672 |
| Aliiroseovarius | 0 | 0 | 0,01774530271 |
| Lentibacter | 0 | 0 | 0,01356993737 |
| Sulfitobacter | 0 | 0 | 0,01043841336 |
| Aurantivirga | 0 | 0 | 0,008350730689 |
| Limimaricola | 0 | 0 | 0,007306889353 |
| Paenibacillus | 0 | 0 | 0,005219206681 |
| Mycoplasma | 0 | 0 | 0,004175365344 |
| Nannocystis | 0 | 0 | 0,003131524008 |
| Paraglaciecola | 0 | 0 | 0,003131524008 |
| Poseidonibacter | 0 | 0 | 0,003131524008 |
| Pseudoalteromonas | 0 | 0 | 0,003131524008 |
