## Supplemental File 2 for "Generating gnotobiotic bivalves: a new method on Manila clam (*Ruditapes philippinarum*)"

**File S2.** Sanger sequence of 16S rRNA gene of *Endozoicomonas* spp. detected in antibiotic-treated clams.

>T3-GF

AGGCCTAACCATGCAAGTCGAGCGGTAACAGGAGGAAGCTTGCTTTCTGCTGACGAGCGGCGGACGGGTGCGTAACACGTAGGAATCTGCCCGGTAGTGGGGGATAGCCCGGAGAAATCCGGATTAATACCGCATACGTCCTAAGGGGGAAAGCAGGGGATCTTCGGACCTTGCGCTATCGGATGAGCCTGCGTCGGATTAGCTAGTTGGTGGGGTAAAGGCCTACCAAGGCCACGATCCGTAGCTGGTCTGAGAGGATGATCAGCCACACTGGGACTGAGACACGGCCCAGACTCCTACGGGAGGCAGCAGTGGGGAATATTGCACAATGGGGGAAACCCTGATGCAGCCATGCCGCGTGTGTGAAGAAGGCTCTAGGGTTGTAAAGCACTTTCAGTAGGGAGGAAAGGGTGGAGGTTAATACCCGTCATCTGTGACGTTACCTACAGAAGAAGCACCGGCTAACTCCGTGCCAGCAGCCGCGGTAATACGGAGGGTGCAAGCGTTAATCGGAATTACTGGGCGTAAAGAGTACGTAGGCGGCTGCCTAAGTTGGATGTGAAAGCCCTGGGCTTAACCTGGGAACTGCATCCAAAACTGGGCGGCTAGAGTGCGGAAGAGGAGTGTGGAATTTCCTGTGTAGCG
