## Supplemental Table 3 for "Generating gnotobiotic bivalves: a new method on Manila clam (*Ruditapes philippinarum*)"

**Table S3.** Relative abundances of all bacterial genera detected in the experimental samples through 16S rRNA sequencing (T1: acclimated clams; T3: antibiotic-treated clams; T4: 1h post-transplant; T5: 6h post-transplant; T6: 22h post-transplant; C: control (non-transplanted) clams; T: transplanted clams).

| **Genus** | **T1** | **T3** | **T4** | | **T5** | | **T6** | |
| --- | --- | --- | --- | --- | --- | --- | --- | --- |
|  |  |  | **C** | **T** | **C** | **T** | **C** | **T** |
| Alteromonas | 0.005548925 | 3.89E-05 | 0.000212 | 0 | 0.000286 | 0 | 0.025429 | 0.001447 |
| Anderseniella | 0.000144076 | 0.000136 | 0 | 0 | 0 | 0 | 0 | 0 |
| Aquimarina | 0 | 0.000278 | 0 | 0 | 0 | 0.000702 | 0.000361 | 0 |
| Blastocatella | 0 | 0 | 0 | 0 | 0.000121 | 0 | 0.000232 | 0 |
| Calorithrix | 2.88E-05 | 9.14E-06 | 0 | 0 | 0 | 0 | 0 | 0 |
| Candidatus Endoecteinascidia | 0 | 0.000849 | 0.003034 | 0 | 0.005225 | 0 | 0.000717 | 0 |
| Candidatus Entotheonella | 0 | 0 | 0.000314 | 0 | 0.000113 | 6.70E-05 | 0.000235 | 0 |
| Candidatus Paenicardinium | 0 | 0.000443 | 0 | 0 | 0 | 0 | 0 | 0 |
| Donghicola | 0.008076506 | 0 | 0 | 0 | 0 | 0 | 0 | 0 |
| Endozoicomonas | 0.91241926 | 0.974694 | 0.972945 | 0.106589 | 0.96075 | 0.314006 | 0.93051 | 0.369613 |
| Filomicrobium | 0 | 0.000133 | 0 | 0 | 0.000148 | 0 | 0 | 0 |
| Flavicella | 0 | 2.50E-05 | 0 | 0 | 0 | 0 | 6.05E-05 | 0 |
| Glaciecola | 0.000496657 | 0.000113 | 0 | 0 | 0.00015 | 0 | 0 | 0 |
| Halomonas | 0.001324689 | 0 | 0 | 0.496065 | 0.000575 | 0.201811 | 0.00093 | 0.195049 |
| Kiloniella | 0.000223535 | 0.000146 | 0 | 0 | 0 | 0 | 0 | 0 |
| Leisingera | 0.004387633 | 0.000306 | 0.000419 | 0 | 0.000316 | 0 | 0.000393 | 0 |
| Lentibacter | 0.002998536 | 0 | 0 | 0 | 0 | 0 | 0.000182 | 0 |
| Limimaricola | 0.000853832 | 0 | 0.000324 | 0 | 2.82E-05 | 0 | 0.000202 | 0 |
| Maribacter | 4.76E-05 | 6.74E-05 | 0.000154 | 0 | 0 | 0 | 0.002062 | 0.000272 |
| Muricauda | 0 | 6.16E-06 | 0 | 0 | 0 | 0 | 0.000316 | 0 |
| Mycoplasma | 0 | 0.00093 | 0.000805 | 0.000593 | 0.004318 | 0 | 0 | 0 |
| NS3a marine group | 0 | 0 | 0 | 0 | 0.000106 | 0 | 5.38E-05 | 0 |
| Nannocystis | 0 | 0 | 0 | 0 | 0.000542 | 0 | 0 | 0 |
| Nitrosomonas | 0.000476655 | 0.000439 | 0.001748 | 0 | 0.000135 | 0 | 0.00071 | 0 |
| Nitrospira | 0.002596031 | 0.000292 | 0.000842 | 0 | 0.00153 | 0 | 0.000433 | 0 |
| Pelagibius | 0 | 4.11E-05 | 0 | 0 | 7.98E-05 | 0 | 0 | 0 |
| Phaeocystidibacter | 0 | 0.000226 | 0 | 0 | 0 | 0 | 0.000454 | 0 |
| Polaribacter | 0.00870711 | 0.000813 | 0 | 0 | 0.000145 | 0 | 0.000609 | 0 |
| Polynucleobacter | 0.000226138 | 0.000195 | 0.000344 | 0 | 0 | 0 | 0.000373 | 0 |
| Pseudoalteromonas | 0 | 0 | 0 | 0 | 0 | 0 | 0 | 0 |
| Pseudomonas | 2.63E-05 | 1.03E-05 | 0 | 0 | 0 | 0 | 0 | 0 |
| Pseudoruegeria | 0.000261844 | 0.000176 | 0.000441 | 0 | 0.000258 | 0 | 0 | 0 |
| Roseobacter clade NAC11-7 lineage | 0.000743875 | 0 | 0 | 0 | 0.000271 | 0 | 0 | 0 |
| Roseovarius | 0.002702253 | 0.000186 | 0 | 0 | 0 | 0 | 0 | 0 |
| Sagittula | 0 | 0 | 0.000112 | 0 | 0 | 0 | 6.39E-05 | 0 |
| Salinimonas | 0.001139021 | 0 | 0 | 0 | 0 | 0 | 0.000222 | 0 |
| Seonamhaeicola | 0 | 0.000556 | 0.000748 | 0 | 0.001321 | 0 | 0 | 0 |
| Shewanella | 0 | 0.000205 | 0.000903 | 0.278434 | 0.002811 | 0.352147 | 0.008458 | 0.296748 |
| Spirochaeta 2 | 0.005510664 | 0.017165 | 0.015708 | 0 | 0.017164 | 0.009771 | 0.024854 | 0.005308 |
| Sulfitobacter | 0.003352373 | 0.001201 | 0.00021 | 9.47E-05 | 0.001139 | 0 | 0.000495 | 0 |
| Tenacibaculum | 0.001082214 | 0 | 0.000592 | 0 | 0 | 0.000363 | 0 | 0 |
| Truepera | 6.90E-05 | 0 | 0 | 0 | 6.70E-05 | 0 | 0 | 0 |
| Umboniibacter | 0.000202334 | 0 | 1.20E-05 | 0 | 0 | 0 | 0 | 0 |
| Vibrio | 0.036354084 | 0.000319 | 0.000132 | 0.118224 | 0.001586 | 0.121133 | 0.001644 | 0.131562 |
| Winogradskyella | 0 | 0 | 0 | 0 | 0.000816 | 0 | 0 | 0 |
